# Conformational Dynamics of Nonenveloped Circovirus Capsid to the Host Cell Receptor

**DOI:** 10.1101/709824

**Authors:** Jiarong Li, Jinyan Gu, Shengnan Wang, Cui Lin, Jianwei Zhou, Jin Lei, Jiyong Zhou

## Abstract

Circovirus, comprising one capsid protein, is the smallest nonenveloped virus and induces lymphopenia. Circovirus can be used to explore the cell adhesion mechanism of nonenveloped viruses. We developed a single-molecule fluorescence resonance energy transfer (smFRET) assay to directly visualize the capsid’s conformational feature. The capsid underwent reversible dynamic transformation between three conformations. The cell surface receptor heparan sulfate (HS) altered the dynamic equilibrium of the capsid to the high-FRET state, revealing the HS binding region. Neutralizing antibodies restricted capsid transition to a low-FRET state, masking the HS binding domain. The lack of positively charged amino acids in the HS binding site reduced cell surface affinity and attenuated virus infectivity via conformational changes. These intrinsic characteristics of the capsid suggested that conformational dynamics is critical for the structural changes occurring upon cell surface receptor binding, supporting a dynamics-based mechanism of receptor binding.

**Importance:** Viral proteins were commom working as ligand to interacte with cell surface glycosaminoglycan receptors to achieve the virus attachment, during which the conformational dynamics of the protein ligand are also vital for the binding properties. In this study, PCV2 capsid and heparin sulfate were used to study the protein conformational dynamics of nonenveloped and icosahedral circovirus capsid during triggering to cell surface receptor. we demonstrated the PCV2 capsid could acts as a dynamic machine, spontaneously adopting multiple conformations with reversible interconversion and intrinsic conformational features could be regulated by glycosaminoglycan receptors and neutralizing antibodies. These increased our understanding of the mechanism by which nonenveloped virus attach to cells.

## INTRODUCTION

Virus attachment to the host cell surface is the initial step in establishing effective infection. The host cell membranes are abundantly decorated with proteoglycans, which comprise a core protein and several types of covalently attached glycosaminoglycan chains (GAGs) that are capable of binding protein ligands (1, 2). GAGs include heparan sulfate (HS), chondroitin sulfate (CS), and keratan sulfate, which are characterized by disaccharide repeating units that form the blocks of the polysaccharides (3–5). Epimerization and terminal sulfation are the most common modification patterns during the processing of GAG chains, and are also vital for the binding properties of the ligands (6, 7). Among these GAGs, HS is utilized by numerous viruses for attachment to cell surfaces, such as the hepatitis C virus (8), human enterovirus 71(9), rabies virus(10), human papillomavirus type 11(11), porcine epidemic diarrhea virus(12), and porcine circovirus type 2 (PCV2)(13). The canonical HS binding motif has been identified as “XBBXBX” or “XBBBXXBX” (“B” represents a basic amino acid and “X” represents a neutral/hydrophobic amino acid) through molecular modeling based on authenticated HS binding protein receptors (14). However, the regulation of virus-ligand conformational features and the mechanism of binding to HS are unknown.

Since 1996, single-molecule Fluorescence Resonance Energy Transfer (smFRET) has been used to analyze replication, transcription, translation, RNA folding, non-canonical DNA dynamics, and protein conformational changes (15–17). The envelope glycoprotein of enveloped viruses is a membrane fusion machine that promotes viral entry into host cells. To date, the conformation dynamics of the envelope glycoproteins of HIV-1 and flu virus have been analyzed. The glycoprotein gp120 of the HIV-1 envelope was shown to have three distinct pre-fusion conformations, whose relative occupancies were remodeled by receptor CD4 and antibody binding (18). In addition, the hemagglutinin of the flu virus envelope was revealed to undergo reversible exchange between the pre-fusion and two intermediate conformations (19). However, for the numerous nonenveloped viruses, the direct visualization and conformational dynamic assessment of the capsid remain unreported.

Circovirus is the smallest nonenveloped icosahedral virus, with a circular, single-stranded genomic DNA, which infects a variety of species ranging from animals to plants with different pathogenicities (20). PCV2, with an unique structural capsid protein, is a representative circovirus that causes significant morbidity and mortality in swine (21–23). The crystal structure of the PCV2 virus-like particle (VLP) revealed that the capsid protein comprises two β-sheets, each containing four antiparallel β-strands, which were labeled as B to I (24). The residues between the β-strands form eight loops, among which the CD and C-terminal loops are exposed on the VLP exterior and can accommodate the insertion of short foreign peptides (25–27). The capsid protein was also reported to have the neutralizing and non-neutralizing epitope sites, and the putative canonical HS binding motif ^98^IRKVKV^103^(13, 28, 29). Recently, Dhindwai et al. also proposed the existence of multiple weak binding sites that interact with HS of the PCV2 VLP surface. However, as the simplest nonenveloped virus, in which the capsid is only composed of one protein, what are conformational features of the circovirus capsid during binding to the cell surface receptor? Does the binding motif move toward to the exterior face of the virus particle to achieve the interaction with the GAG molecules? What is the key factor of the motif and the GAGs that facilitate the interaction by adjusting of the capsid’s conformational behavior? In the present study we aimed to answer some of these using the PCV2 Capsid and HS as a model to explore the conformational dynamics of nonenveloped viruse invasion of cells.

In the present study, we developed an smFRET-imaging assay to detect the conformational dynamics of the virus capsid protein (Figure 1a). Our results revealed that the Capsid worked in a dynamic manner, undergoing spontaneous and reversible transitions between three distinct conformations. The binding of the GaGs and that of neutralizing antibodies shifted the dynamic equilibrium associated with the virus binding to the host cells. Both the negatively and positively charged distribution of the receptor and the binding motif peptides played a critical role in the capsid’s conformational dynamics, and ultimately influenced the affinity for the cell surface receptor. The allostery of the capsid occurs during direct interactions with the GAG molecules might provide a strategy by which the circovirus strengthen its attachment to the host cells, ultimately resulting in infection.

**Figure 1.**
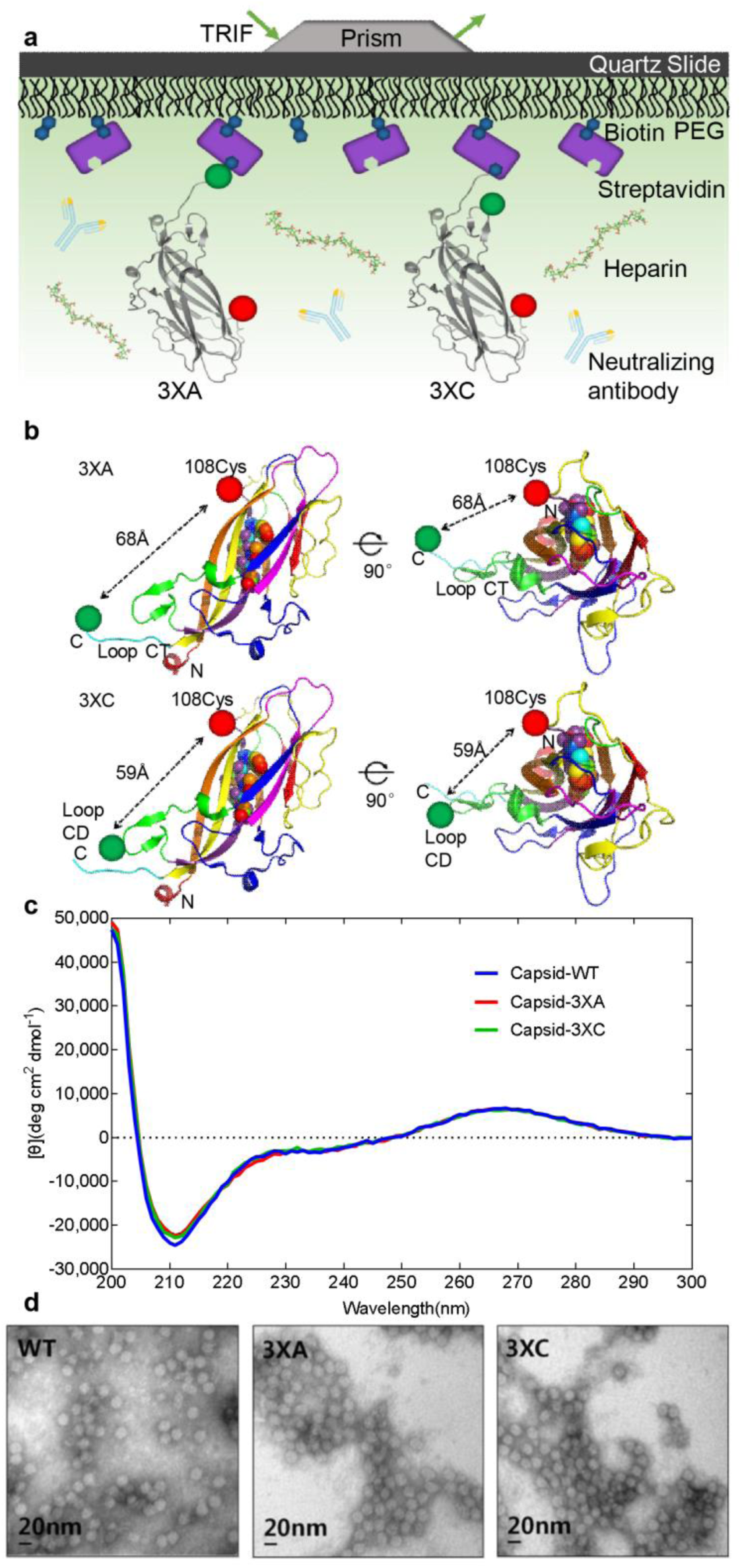
Single-molecule FRET Method for Direct Capsid Visualization. (a) Schematic of the smFRET-imaging assay. Capsid 3XA or 3XC protein monomers containing a single pair of dually labeled fluorophores were immobilized on quartz microscope slides, and imaged using TIRF microscopy at room temperature. (b) Structural molecular models of Cy5/Alexa 547-labeled Capsid. Fluorophores Cy5 (red ball) and Alexa 547 (green ball) were placed at the 108cysteine residue and at C-terminal loop (3XA) or at the CD loop (3XC), respectively. These labeled models reflected the conformation transitions by their changes in FRET efficiency. The putative HS binding site within the capsid is represented as a sphere. All the structures were adapted from PDB accession ID:3R0R. (c) Circular dichroism assay of the secondary structure of soluble WT, 3XA, and 3XC capsids. The spectrum curve of the samples was acquired from the ellipticity data measured at a wavelength of 200¬–300 nm. (d) Transmission electron microscopy imaging of PCV2 virus-like particles. Capsid-WT, Capsid-3XA and Capsid-3XC assembled into a hexagonal particle with a diameter of 17 nm.

## RESULTS

### Site-specific Attachment of Fluorophores to the Capsid Protein

To develope an smFRET-imaging assay to visualize the conformational dynamics of the PCV2 capsid (Figure 1a), we attached the Cy5 fluorophore (acceptor fluorophore) at position ^108^Cysteine in the DE loop of the capsid(Figure S1a)(30). To install the donor fluorophore, we inserted the A1 Tag peptide (GDSLDMLEWSLM) at position ^80^Leucine of the CD Loop or at position ^233^Proline of the C-terminal Loop, which are relatively non-conserved regions of capsid. Thus, the Alexa 547 fluorophore was labeled using holo-acyl carrier protein synthetase (ACPs), which catalyzed serine hydroxylation of the A1 peptide and the 4’-phosphopantetheine moiety of CoA to form a phosphodiester bond (31). The Avi-peptides were attached at the C-terminus of the capsid to accomplish biotinylation processing via the BirA catalyzation (Figure S1a) (32). According to previously reported structure of the capsid, these two designed models, termed Capsid 3XA (3XA) and Capsid 3XC (3XC), were used to demonstrate that the distances between the donor and acceptor fluorophores was 68 Å (3XA) and 59 Å (3XC) (Figure 1b), which satisfied the demand of the smFRET experiment (30). Western blotting and typhoon assays confirmed Cy5 was co-localized with Alexa 547, demonstrating that 3XA and 3XC were specifically labeled with fluorophore pair and biotin (Figure S1b).

After optimization of the labeling reaction *in vitro*, the labeling efficiencies of Alexa 547 and Cy5 were approximately 45% and 71%, respectively, indicating that approximately 32% of molecules would be suitable for smFRET analysis using total internal reflection fluorescence (TIRF) microscopy. To assess whether the secondary structure of the capsid was disturbed by the insertion of exogenous peptides, we performed a circular dichroism (CD) assay of soluble wild-type (WT) capsid, 3XA, and 3XC over the spectrum of 200–300 nm. The CD spectrum curves of 3XA and 3XC were identical to that of the WT capsid (Figure 1c), indicating that the insertion of exogenous peptides in 3XA and 3XC had a negligible effect on the protein secondary structure. Using transmission electron microscopy, we also observed that the 3XA and 3XC monomers could self-assembly into a hexagonal virus-like particle with a diameter of almost 17 nm (Figure 1d), similar to the PCV2 virion, indicating that the intermolecular interaction of the capsid monomers was still viable. Taken together, 3XA and 3XC, with short amino acid modifications, retained the intrinsic structural features similar to the WT capsid, and could also be imaged using TIRF microscopy.

### Capsids Spontaneously Interconvert between Three Distinct Conformations

To disperse the capsid into monomer and minimize the disturbance of oligomerization to imaging on the TRIF microscopy, we treated 3XA and 3XC with imaging buffer containing different concentrations of NaCl for 30 min. The capsids treated with 1500 mM NaCl were immobilized on the covered quartz microscope slide in a single molecule state (Figure S1c). We first visualized the conformational landscape of 3XA and 3XC, the smFRET trajectories of 3XA displayed a spontaneous reversible model with fluctuations between three conformations characterized by distinct FRET values of approximately 0.29, 0.48, and 0.68. Similarly, 3XC exhibited a conformational distribution with distinct FRET values of approximately 0.39, 0.59, and 0.78 (Fig. 2a). Both the smFRET trajectories displayed a predominant low-FRET state with a low frequency of transitions to the intermediate- and high-FRET states, corresponding to 77% and 79% low-FRET state occupancies in the histograms of the 3XA and 3XC FRET trajectories, respectively (Figure 2b). The FRET values of the low-FRET state were consistent with the 68Å and 59Å inter-fluorophore distance predicted based on the structure of the of 3XA and 3XC. Moreover, a direct transition between the low- and high-FRET states of the capsid was hardly observed (Figure 2a). Collectively, these data demonstrated that the low-FRET state was the predominant conformation for 3XA and 3XC, and the intermediate FRET state could be essential for the transition between low- and high-FRET states.

**Figure 2.**
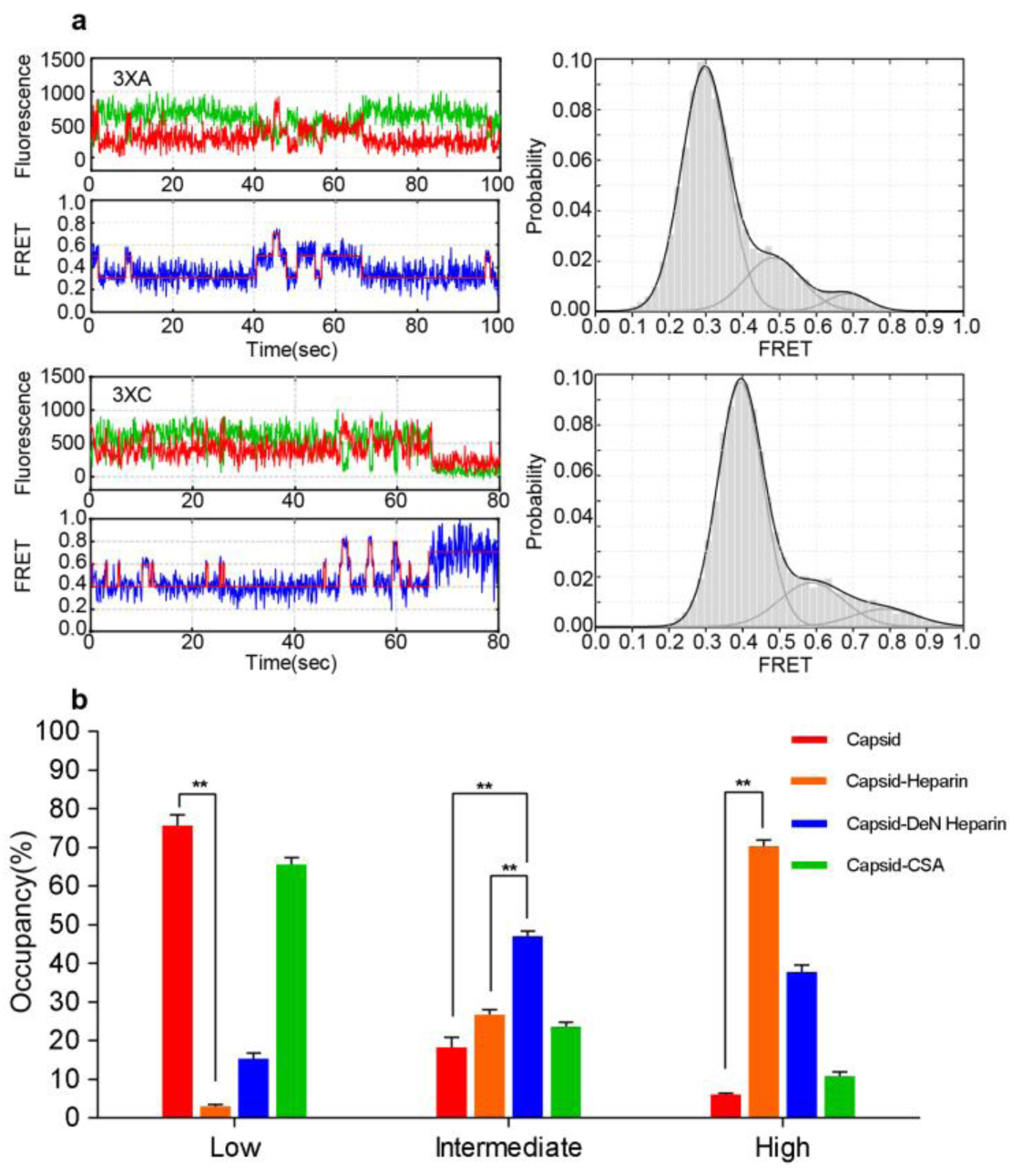
The Multiple Conformational Distributions of the Capsid Protein. (a) The landscapes of the capsid’s conformational features. Representative fluorescence intensities of the donor and acceptor are shown as green and red, respectively, in the time trace; the original and idealized FRET values are shown as blue and red, respectively, in the FRET trajectories (left). The distinct FRET values of the main states fitted using HMM were approximately 0.29, 0.48, 0.68 (3XA) and 0.39, 0.59, 0.78 (3XC), corresponding to population FRET histograms fitted with three Gaussian distributions overlaid (black) (right). (b) Occupancy of the smFRET state in Figures 2A and 3A-C. Student’s t test was performed. *, *P* < 0.05; **, *P* < 0.01.

### Binding to Heparan Sulfate Alters the Conformational features of the Capsid

Heparan sulfate (HS) is the cell surface receptor for PCV2 attachment (13). The negative charges on the GAGs are acquired through N- and O-sulfation of the carbohydrate moieties and are crucial for their interactions (33). To determine if the conformational equilibrium of the capsid changed during binding to cell receptor HS, we performed smFRET analysis of 3XA and 3XC with the addition of heparin, an analog of heparan sulfate with a similar structure to HS’s sulfated domain. Compared with the smFRET data of free 3XA and 3XC, the trajectories of capsids with heparin exhibited a high-FRET-preponderant model and intermittently dropped to an intermediate state or low state for both 3XA and 3XC. The occupancy of the high-FRET state was significantly elevated to ∼70% from ∼6.3%, and the occupancy of the low-FRET state significantly decreased to ∼4% from ∼77% (Figures 2, 3A and S2A; *P* < 0.001), demonstrating that the high-FRET state is the preponderant state of the smFRET trajectory for the capsid-heparin binding complex.

Interestingly, in contrast to the capsid-heparin complexes, the mixture of capsid and De-N-sulfated acetylated heparin (DeN heparin, a highly sulfated HS analog without N-sulfation used to represent the incompletely negative charged HS) exhibited an marked increase in the intermediate FRET state (47%) and high-FRET state (37.7%), as well as a decrease in the low-FRET state (Figures 2B, 3B and S2B). However, as a negative control, a mixture of capsid and chondroitin sulfate A (CSA, another member of the cell surface glycosaminoglycans but does not act as a binding receptor during the PCV2 infection) exhibited a similar conformational distribution to the free capsid (Figures 2, 3c and S2c). These findings demonstrated that the binding of negatively charged heparin altered the conformational features of the capsid and that the high-FRET state of the capsid is the main conformation during the HS interaction.

To verify whether the changes in capsid conformation affected the affinity to host cells, we performed GAGs competitive binding experiments with PK-15 and 3D4/31 cells. The results of immunoblotting and quantitative real-time PCR (qPCR) assays showed that both the level of bound VLPs and attached PCV2 particles to host PK15 and 3D4/31 cells in the presence of added soluble heparin decreased significantly compared with those of free VLPs and PCV2; however, the addition of DeN heparin or CSA had no significant effect (Figure 3d-e and S2d-e). In addition, in the flow cytometry assay, cell adhesion of VLP and PCV2 in the presence added GAGs showed that for PK15 cells, 70.3% bound VLP and 30.4% bound PCV2, which was an obvious decrease compared with free VLP and PCV2, VLP and PCV2 with added DeN heparin, and VLP and PCV2 with added CSA (Figure 3f-g). Surprisingly, the addition of DeN heparin to VLP and PCV2 showed almost similar occupancies to those of VLP and PCV2 attached to cells compared with the CSA addition as a negative control. This result demonstrated that heparin without a negative charge acquired from N-sulfation is a nonfunctional HS analog for binding VLP and PCV2 virions. The results obtained using 3D4/31 cells were similar to those gained using PK15 cells (Figure S2f-g). In summary, soluble heparin can bind to the capsid and promote the high-FRET state, simultaneously block the binding of the capsid to the host cells.

**Figure 3.**
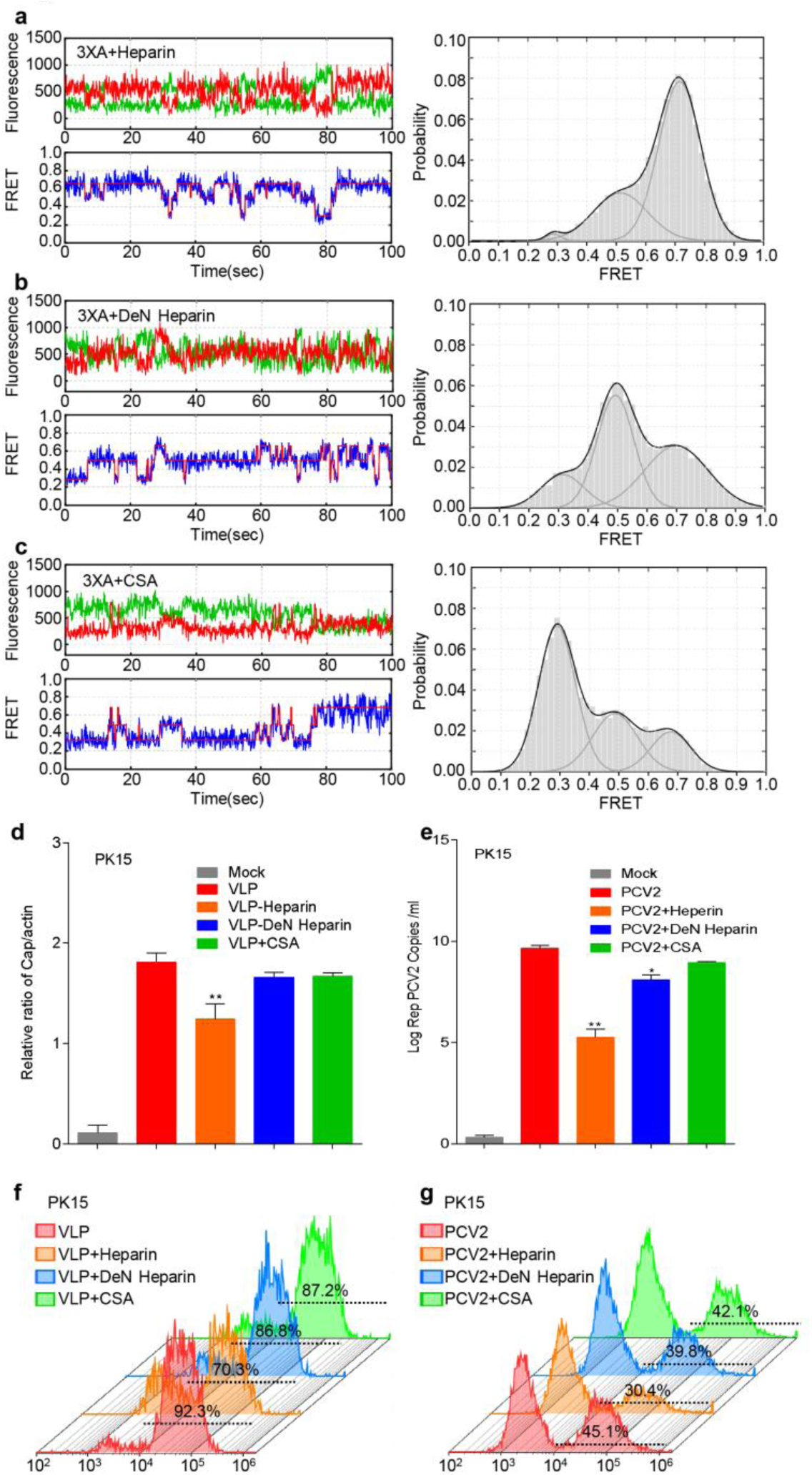
Interaction with Heparin Regulates the Capsid Conformational landscape. (a-c) The histogram distribution and representative fluorescence time trace of the capsid protein’s interaction with GAGs. a, Capsid mixed with heparin. b, Capsid mixed with De-N-sulfated acetylated heparin. c, Capsid mixed with chondroitin sulfate A. The concentration of GAGs was fixed at 2500 μg/ml. (d-g) GAGs competitive binding experiments to host cells. Equal amounts of capsid-VLPs or PCV2 virions were preincubated with heparin, De-N-sulfated acetylated heparin, and chondroitin sulfate A at 37 °C for 90 min, resp ectively. The mixtures were then seeded onto PK-15 cells for 60 min at 4 °C. (d), the total amount of capsid protein bound to the cell surface was analyzed using SDS-PAGE and western blotting. (e), the number of copies of attached PCV2 virions was quantified using qPCR. (f), capsid-VLPs bound to PK15 cells as assessed using flow cytometry. (g), PCV2 attached to PK15 cells as assessed using flow cytometry.

### Capsid-targeting Neutralizing Antibodies Facilitate Stable Low-FRET Conformations

To investigate whether anti-capsid neutralizing monoclonal antibodies (mAbs) could block the virus binding to the cell surface by modifying the conformational transitions of the capsid protein, we incubated 3XA and 3XC with the mAbs 3F6, 5E11, 6H9, and 8A12 before smFRET imaging (29). Interestingly, the neutralizing mAb 3F6 induced an absolute occupancy of the low-FRET state (94%) of the capsid and resulted in the almost compete disappearance of the intermediate-FRET and high-FRET states compared with the trajectories of the antibody-free capsid (Figures 4a and 2a). Another neutralizing mAb, 6H9, displayed similar effects to 3F6 (Figure S3a). Interestingly, although transition to the intermediate-FRET state and high-FRET state were obviously inhibited, they were not entirely abolished, implying that the mobility of the capsid was maintained. In contrast, the conformation of the capsid after treatment with non-neutralizing mAbs 5E11 and 8A12 was similar to that of the antibody-free Capsid, with only a slight increase in the intermediate-FRET state, demonstrating the negligible effect of nonneutralizing mAbs in altering the antigen’s conformation (Figures 4b, S3b and 2a). Based on these results, we concluded that neutralizing antibodies could restrict the conformation of the capsid to the low-FRET state by inducing an almost complete conversion of the capsid’s conformation from the intermediate- and high-FRET states to the low-FRET state without impairing the mobility of the antigen.

**Figure 4.**
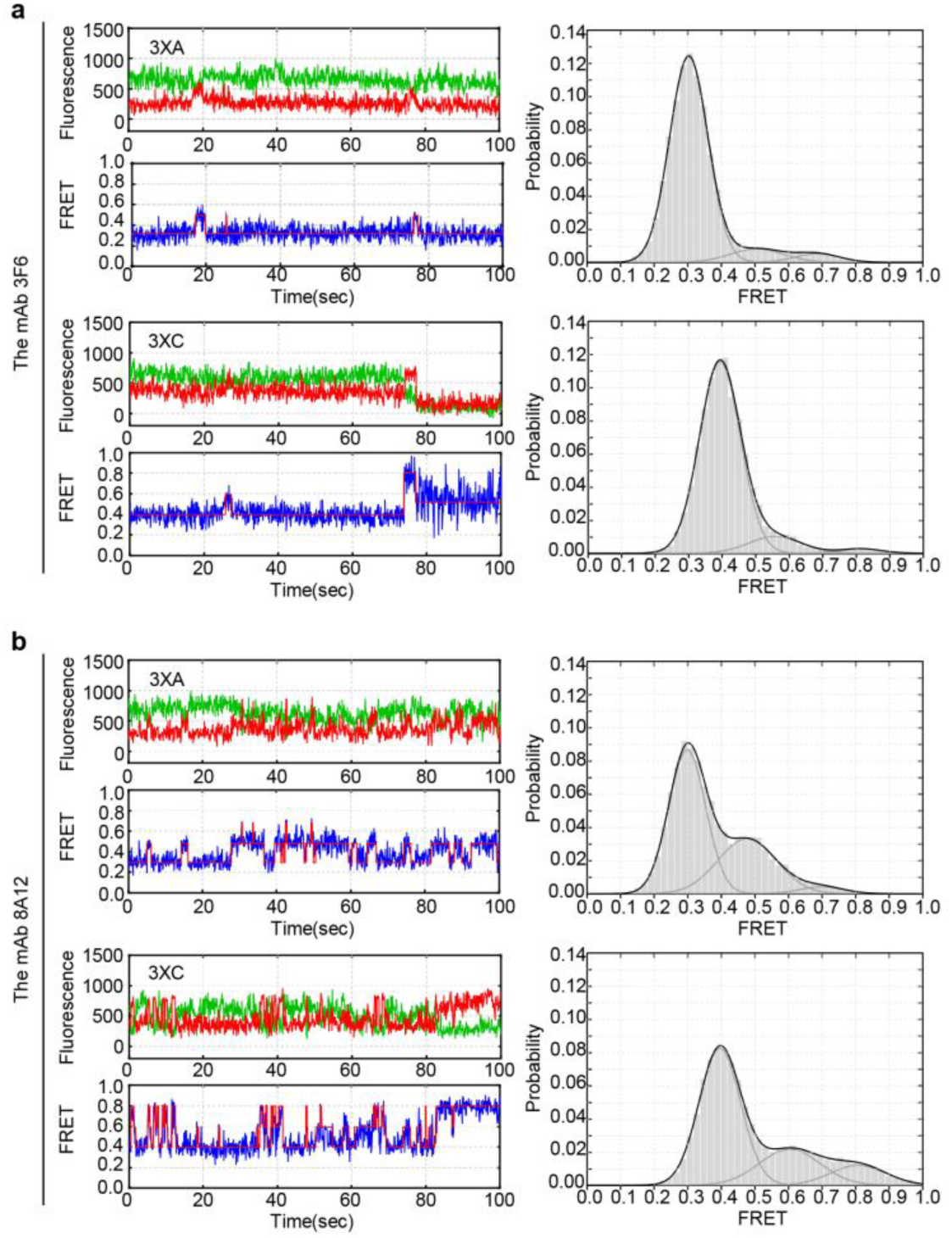
Neutralizing Antibodies Restrict smFRET Conformations of the Capsid Protein. Representative fluorescence time trace and histogram distribution of the anti-Capsid mAbs reaction with capsids 3XA and 3XC. (a) Capsid binding to the neutralizing mAb 3F6 reveals an absolute predominance of the low-FRET state. (b) Capsid binding to the non-neutralizing mAb 8A12 reveals a similar conformational feature to free capsids.

### The Positively Charged Residues ^99^R^100^K within the HS Binding Motif of the Capsid Enhance the High-FRET State During Interaction with Heparan Sulfate

The motif ^98^IRKVKV^103^ within the capsid protein is predicted to be the binding site of HS, and is located in the interior of the three-dimensional crystal structure of the capsid (13). The interaction between the GAGs and a protein mainly depended on the positively charged properties of the amino acids in the binding motif (34). Amino acid sequence alignment revealed that the predicted HS binding motif ^98^IRKVKV^103^ is highly conserved among different subtypes of PCV2 (Figure S4a). To investigate the contribution of positively charged amino acids to conformational changes during the interaction with HS, we performed ^99^ arginine (R) to alanine (A) single-point mutation (SM), and ^99^R^100^lysine (K) to A double-point mutations (DM) within the HS binding site of the capsid (Figure S4b). We subjected the dual-labeled Capsid-SM and Capsid-DM to smFRET analysis, which displayed similar conformational dynamics to Capsid-WT (Figures 2a, 5a-b and S4c-d), such as the interconversion between the three intrinsic states and a predominant occupancy of the low-FRET state with occasional transition to intermediate- and high-states. This result implied that the positively charged residues ^99^R^110^K within the HS binding motif did not directly regulate capsid conformational dynamics in the absence of receptor.

To further verify the role of the positively charged amino acids during the capsid’s interaction with HS, we observe the smFRET landscapes of Capsid-SM and Capsid-DM in the presence of heparin. The trajectories of Capsid-SM and Capsid-DM still displayed the three independent states; however, the trajectories were shifted toward increased intermediate- and high-FRET states (Figure 5c-d and S4e-f). The data acquired from Capsid-SM in the presence of heparin demonstrated that the occupancy of the high-FRET was ∼54%, which was significantly lower than that for Capsid-WT in the presence of heparin (∼77%), and the occupancies of intermediate and low FRET states both showed various degrees of upregulation (Figures 5c, 3a, S2a and S4e). This tendency was more obvious in the mutant lacked two positively charged amino acids in the binding motif. The trajectories of Capsid-DM in the presence of heparin displayed more frequent transitions to the intermediate and low-FRET states compared with Capsid-SM and Capsid-WT (Figures 5d, 3a, S2a and S4f), resulting in a further reduction of high-FRET state occupancy (∼38%), accompanied by a slight increase in the intermediate-FRET state occupancy and increased low-FRET state occupancy. This data indicated that replacement of the positive charge amino acids within the HS binding motif decreased the occupancy of the high-FRET state significantly during the interaction with heparin. We further delineated the relationship between the numbers of positively charged amino acids and the occupancy fluctuation of each state. Figure 5e shows that decreasing high-FRET occupancy correlated positively with the decreasing number of positively charged amino acids. By contrast, the occupancies of intermediate- and low-FRET states correlated negatively with the number of positively charged amino acids (Figure 5e). Taken together, these findings indicated that the positively charged residues ^99^R^100^K within the HS binding motif of the capsid associating with the maintaining of high-FRET state during the interaction with HS.

**Figure 5.**
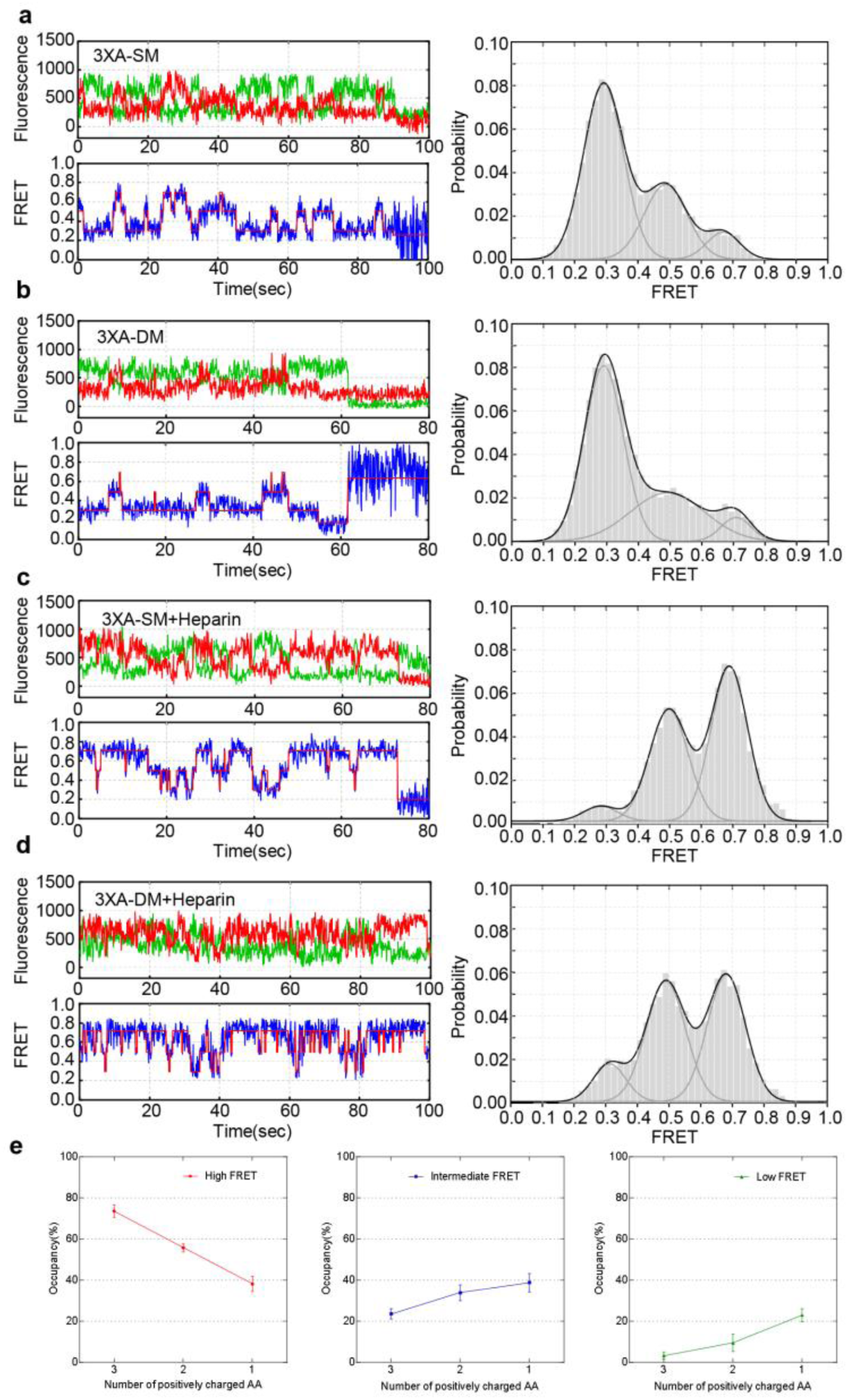
Deficiency in Positively Charged Amino Acids in the Heparan Sulfate Binding Site Alters the smFRET Conformational Feature of the Capsid Under the Effect of Heparin. (a-b) Conformational features of capsids deficient in positively charged amino acids. The smFRET trajectories of Capsid-SM and Capsid-DM showed the conformational feature of interconversion between the three intrinsic states. The histogram shows the distribution of the distinct states, similar to the landscape of Capsid-WT. (c-d) Representative fluorescence trajectories and histogram distributions of Capsid-SM and Capsid-DM mixed with heparin. (e) Curve of each state occupancy in relation to positively charged amino acid numbers within the binding motif based on the histogram data in Figure 5a-d.

### Residues ^99^R^100^K of the HS Binding Site are Critical for The Affinity of the Capsid to the Host Cell

To further investigate the contribution of positively charged amino acid residues within the HS binding motif of the capsid to receptor binding, we performed binding force and cell adhesion tests of Capsid-SM and Capsid-DM. As expected, the amount of Capsid-SM and Capsid-DM that bound to the HiTrap™ Heparin-Sepharose HP Column was significant decreased compared with that of the WT capsid (Figure 6a). Simultaneously, the dissociation constants (Kd) of SM (821 ± 18.1 μM) and DM (883 ± 15.8 μM) displayed an obvious increase compared with that of WT Capsid (252 ± 18.7 μM) measured by microscale thermophoresis (Figure 6b). This indicated that the lack of ^99^R^100^K within the HS binding motif ^98^IRKVKV^103^ would critically impair the affinity of the capsid for heparin.

**Figure 6.**
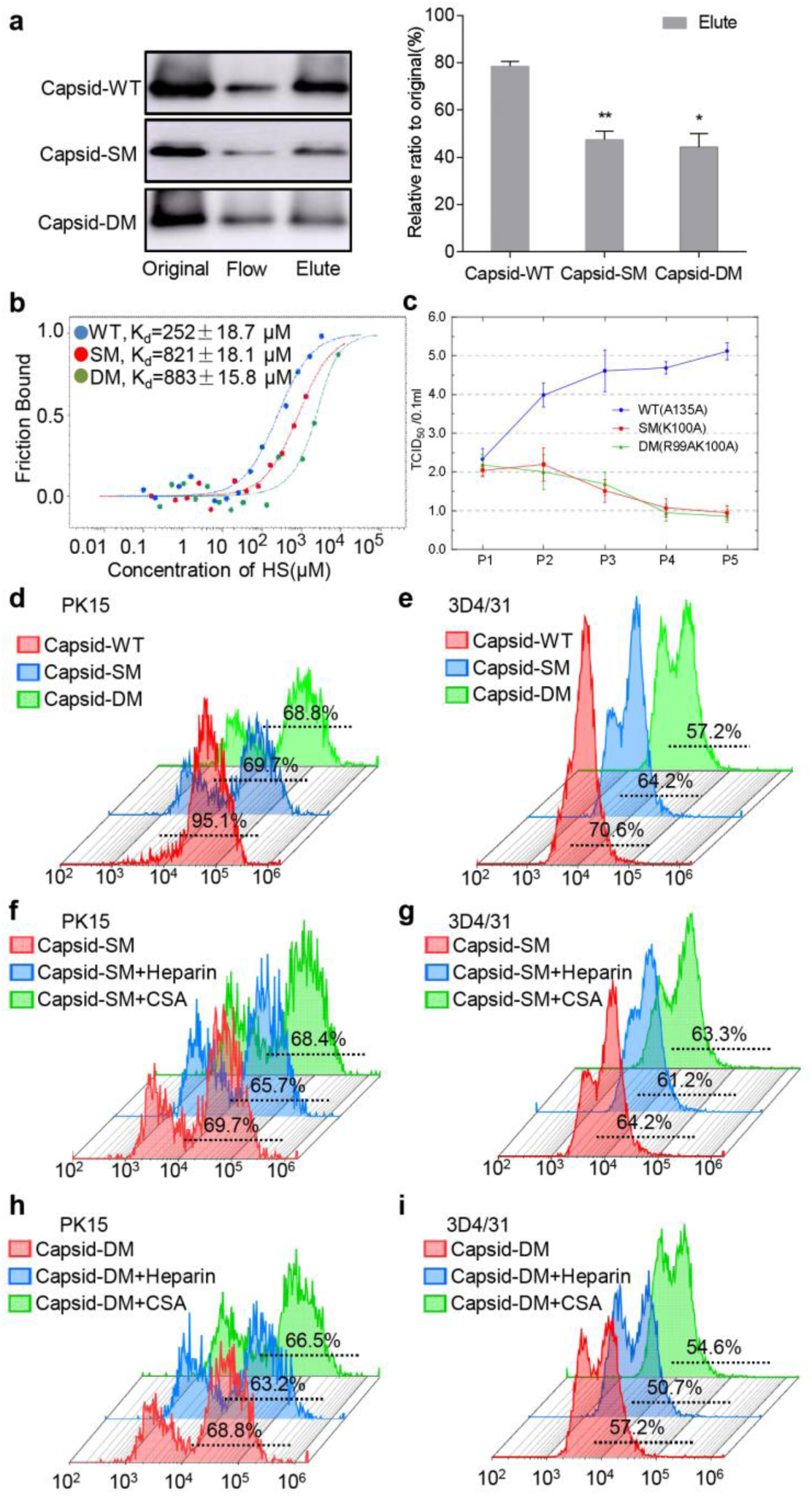
The Charged Amino Acids within the Binding Site Regulate the Affinity of Capsid & Virion for the Host Cells. (a) Relative affinity of the WT, SM, and DM capsids to heparin was measured using HiTrap™ Heparin-Sepharose HP Column chromatography. The ratio of eluted to original amount was calculated to evaluate the relative affinity to Heparin-Sepharose of each sample. (b) The dissociation constant (K_d_) between the capsid and heparin was measured using microscale thermophoresis. The Cy5 labeled WT, SM, and DM capsids were mixed at equal amounts with various concentration of heparin for 15 min at room temperature, and the K_d_ values were determined using microscale thermophoresis. (c) Infectivity decrease of rescued viruses with deficient in positively charged amino acids in the binding motif. TCID50 titration was performed according to the Reed-Muench method. (d-i) Capsid-bound cells measured by flow cytometry. The PK15 and 3D4/31 cells were incubated with Capsid-WT, Capsid-SM, Capsid-DM, or mixtures of Capsid-SM/Heparin, Capsid-DM/Heparin, Capsid-SM/CSA, and Capsid-DM/CSA, separately. The resultant cells were assessed by flow cytometry. (d), Capsid positive PK15 cells; (e), Capsid positive 3D4/31 cells; (f), Capsid-SM positive PK15 cells; (g), Capsid-SM positive 3D4/31 cells; (h), Capsid-DM positive PK15 cells; and (i), Capsid-DM positive 3D4/31 cells.

Given that the affinity of the capsid for heparin was weakened by partial replacement of positively charged amino acids, we next asked whether the lack of these residues would disturb the virus lifecycle. Therefore, we performed virus rescue of PCV2 with the SM and DM mutations. The self-cyclized PCV2 genomes with the ^99^A or ^99^A^100^A mutation of the capsid gene were transfected into PK-15 cells to rescue the virus, which was serially passaged to detect infectivity using the TCID50 assay (35). The one-step growth curve indicated the genomes with the SM or DM mutations could be rescued; however, the infectivity of the rescued virus was weakened compared with that of the WT virus (Figure 6c), demonstrating that the replacement of these positive charged residues within the HS binding site of the capsid could attenuated the replication ability of PCV2. To further validate whether the weakened replication ability of PCV2 particles with SM or DM involves cell attachment, we attempted single cell adhesion assays of SM- and DM-Capsid proteins using flow cytometry. It showed the number of adhesive cells presenting Capsid-SM or Capsid-DM molecules was significant decreased compared with that of the WT Capsid (Figures 6d and 6e). However, the number of adhesive cells in the presence of HS was not significantly different to that of free Capsid-SM or Capsid-DM (figures 6f, 6g, 6h and 6i), suggesting that the positive charge deficiencies in the HS binding motif caused the capsid non-sensitive to HS and reduced the cell binding ability. Thus, these *in vivo* and *in vitro* experiments demonstrated that the positively charged residues ^99^R^100^K within the HS binding motif of the capsid are critical to enhance virus attachment to the host cell surface and to regulate virus infectivity.

## DISCUSSION

Heparan sulfate, a GAG present on the cell membrane, was identified as the cell surface receptor for various proteins and enveloped and nonenveloped viruses (36–38). In the present study, PCV2 was used as a model nonenveloped virus, who is the smallest DNA virus with capsid comprises one unique capsid protein, and used HS as the receptor for attachment to host cells (13, 24). The direct observation of the capsid conformational changes for nonenveloped viruses has been not yet been achieved because of the lack of methodology to probe such dynamics on a relevant timescale. In the present study, the usage of smFRET imaging provided the first real-time visualization of the conformational dynamics of the capsid. The results of the present study indicated that the Capsid may be regarded as a dynamic machine, undergoing spontaneous, reversible fluctuations between multiple conformations during attachment to the cell surface. Our results suggested that receptor binding regulates the intrinsic conformational dynamics of the capsid, allowing the virus to invade the host cells.

### The Multiple Conformations of the Capsid with Spontaneous Reversible Interconversion

The crystal structure of the PCV2 VLP revealed that the HS-binding motif ^98^IRKVKV^103^ was deeply hidden in the interior of the VLP(13, 24). In the present study, Cy5 and A547 were labeled at ^108^cysteine (Cys) near the HS-binding site and at the CD loop or C-terminal loop far from the HS-binding site of the capsid (Figure S1a). The smFRET approach presented a landscape in which the capsid spontaneously transited among at least three conformations, represented by the low-, intermediate- and high-FRET states and the low-FRET state of the capsid, corresponds to the previously described crystal structure of PCV2 VLP (24). However, the current three-dimensional structure of the VLP does not depict the conformations corresponding to the intermediate- and high-FRET states, which we captured using the smFRET assay. The capsid displayed a predominantly low-FRET landscape, with some transitions to the intermediate- or high-FRET states (Figure 2a). Notably, the structural proteins of enveloped viruses, such as glycoprotein gp120 subunit of the HIV-1 envelope trimmers (18) and hemagglutinin of influenza virus (19), show similar dynamic interconversions between multiple distinct conformations. Thus, for the first time, we demonstrated that the capsid of a nonenveloped circovirus could spontaneously and reversible interconvert among multiple conformations. Technical limitations meant that we could not detect the conformational dynamics of capsid in real-time in the form of icosahedral virus particles or VLPs; therefore, unfortunately, we could not verify that the conformational features of the Capsid monomer were recapitulated in the assembled virus particle.

### GAGs and Neutralizing Antibodies Alter the Capsid’s Conformational feature

Heparan sulfate acts as the principal GAG receptor for the attachment of PCV2 capsid, which is partially mediated by electrostatic binding between the negatively charged chains of HS and basic amino acids within the target proteins(1, 39). Indeed, our binding experiments emphasized the superiority of HS over CSA and further demonstrated the requirement for N-sulfation of heparin for interaction with the capsid. The dynamic equilibrium of the capsid was entirely rearranged by negatively charged heparin, ultimately triggering the emergence of the high-FRET state (Figures 3a and S2a). Notably, the negative charge-deficient heparan analog DeN heparin could induce transition from the low-FRET state to the intermediate-FRET sate, but was insufficient to induce the high FRET state (Figures 3b and S2b). However, the competitive binding assay showed that DeN heparin was incapable of binding to the capsid protein or the virus (Figures 3d-g), indicating that the intermediate-FRET state was not an effective conformation for receptor interaction. Therefore, we hypothesized that the high-FRET state is indispensable to the efficient interaction between the capsid and the GAG receptor, and the hidden HS binding site might be exposed on the virion surface when negatively charged heparan sulfate is encountered (Figures 7). However, the current technique could not distinguish if the capsid conformational transitions occurred after the receptor binding to sustain the interaction or if the conformational transitions were induced by the receptor molecule before the binding event (40).

**Figure 7.**
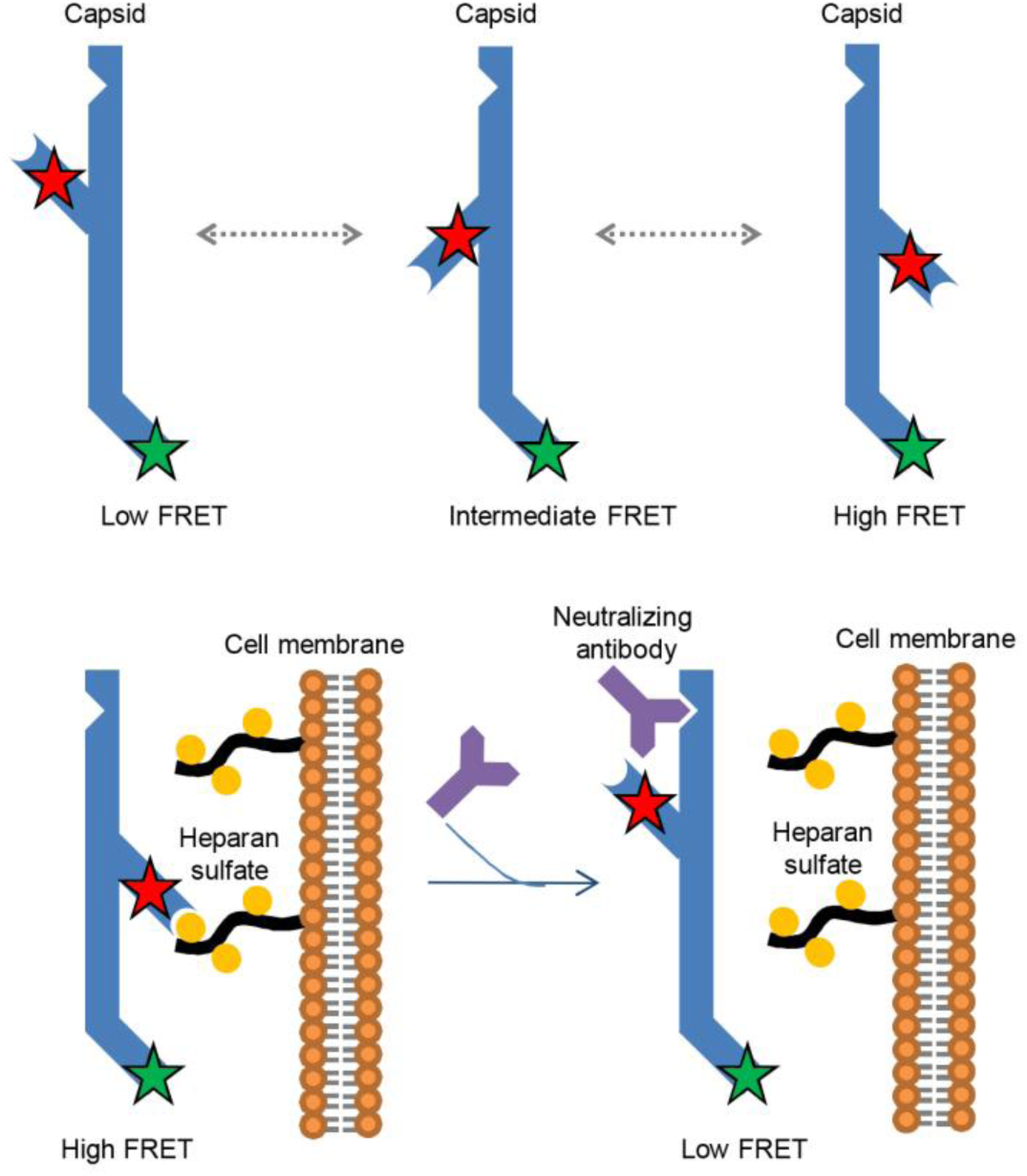
Model of Capsid Conformational changes associated with attachment to the cell surface. Interaction with the receptor HS promotes the preponderance of the high-FRET state and exposes the hidden HS binding site. By contrast, interaction with the anti-Capsid neutralizing mAb results in prevention of the conformational transition and restriction to the low-FRET state, thereby blocking exposure of the HS binding site and preventing the interaction with the cell receptor heparan sulfate.

In contrast to the effect of HS, the capsid showed a tendency to stabilize in the low-FRET state in the presence of the anti-capsid neutralizing mAbs (Figures 4a and S3a), indicating that the neutralizing antibodies prevented the capsid from attaching to the cell surface by restricting the conformational dynamics and consequently disrupting the interaction between the capsid and HS. Interestingly, the antigenic epitope of the capsid recognized by neutralizing mAb 3F6 is ^156^YHSRYFT^162^(29), but not the HS-binding site. So we hypothesized that the interaction with the neutralizing mAb prevented the conformational transitions and restricted the capsid to the low-FRET state, thereby blocking the allosteric effects that expose the HS binding site to allow interaction with cell receptor (Figures 7). This could be considered as indirect evidence for the significance of the high-FRET state during the capsid-cell receptor interaction.

### Charge Distribution of the Binding Motif Dictated the Effective Receptor-ligand Binding

The key factors that produce a hydrophilic pocket producing a complementary structure for the heparin-protein interaction are clusters of basic amino acids, especially arginine and lysine. The mutants with 99R or 100K displayed the markedly attenuated affinity for heparin and reduced the binding capacity to the cell (Figures 6), subsequently impaired the infectivity of the rescued viruses. Likewise, the occurrence of the high-FRET state also decreased markedly as the number of positively charged amino acids in the binding motif decreased. Nonetheless, the conformational transition could still occur in the mutants in the presence of heparin (Figure 5c-d and S3e-f) and the binding of the mutants was not completely abolished, indicating that the interaction could be supported by the remaining positively charged amino acid(s) within the motif or by the existence of other binding sites, such as the weak HS binding sites on the surface of the PCV2 VLP.

### Biological significance of Conformational Feature Regulation During Virus Attachment

The high-FRET state was revealed to be the crucial configuration during the Capsid-HS interaction for virus binding. This state reflected a distinct conformation in which the hidden HS binding site is vicinal to the exterior area of the virus particle. Besides, the existence of the intermediate-FRET state was especially conspicuous during the smFRET assays, but ultimately could not accomplish the receptor-ligand interaction. The existence and maintenance of the intermediate-FRET state could be quite meaningful and important for the biochemical events of the virus. It might prevent the conformational transition to the ultimate structure occurring too quickly when induced by an incomplete interforce and other biochemical attractive forces, thus minimizing meaningless binding to inappropriate receptors.

## Materials and Methods

### Protein Purification and Fluorophore Labeling

Purified proteins were dissolved in labeling buffer (50 mM HEPES, 10 mM MgCl2, 10 mM CaCl2, 150 mM NaCl, pH 7.5), preincubated with a 50-fold molar excess of Tris (2-carboxyethyl) phosphine(T2556, Thermo Fisher) to reduce inter-molecular disulfide bond for 30 min at room temperature, then mixed with 10-fold molar excess CoA-Alexa 547 (S9349S, New England Biolabs) and 5 µM ACPS at 37 °C for 90 min, followed by mixing with 20-fold molar excess of Cy5-maleimide(PA25031, GE healthcare) at room temperature overnight (25, 34). Dual-labeled capsid proteins were separated using a PD Mini-Trap G-25 desalting column (GE healthcare) with imaging buffer (50 mM Tris, 150 mM NaCl, 1 mM Trolox(238813, Sigma-Aldrich), 0.8% glucose (w/v)(G8270, Sigma-Aldrich), 0.8% glucose oxidase (w/v)(G3660, Sigma-Aldrich), 0.8% catalase (w/v)(SRE0041, Sigma-Aldrich), pH 7.5). The labeling efficiency was calculated from protein concentration measured using BCA and fluorescence concentration estimated based on Lambert-Beer equation with absorbance at 550 & 630 nm. Finaly sample was stored at −80 °C until use.

### Circular Dichroism Spectrum Analysis

Purified Capsids were measured using a Chirascan™-Circular Dichroism (CD) Spectrometer (Applied Photophysics) with a path length of 0.1 mm. The CD spectra of the capsids were acquired between 200 and 300 nm with a 1-nm increment, and the spectrum of the free buffer solution was subtracted as background.

### Self-assembly of PCV2 Virus-like Particles

Purified capsids were dialyzed in assembly buffer (NaH2PO4 0.1 M, Na2HPO4 0.1 M, Imidazole 20 mM, Tris base 0.01 M, NaCl 0.15 M, KCl 0.05 M, MgCl 0.002 M, ammonium 0.1 M, glycerol 5%, Triton-X100 0.5%, β-mercaptoethanol 5 mM, and PMSF 0.1 mM, PH-7.6) at 4 °C overnight, then subjected to an ÄKTA Purifier UPC 100 system (GE Healthcare) equipped with a prepacked HiPrep™ 16/60 Sephacryl™ S-300 HR column (GE Healthcare). The formation of PCV2 VLPs was observed using transmission electron microscopy (H-7650 TEM, Hitachi-Science &Technology, Japan).

### Single-molecule FRET Assay

The smFRET experiment was performed based on a custom-built prism-based total internal reflection fluorescence (TIRF) microscopy method (Olympus IX81 microscope, Photometric Evolve 512 EMCCD, 50 mW, Coherent 532 nm solid-state laser). 10 μM of samples was immobilized on a PEG(900393, Sigma-Aldrich)-passivated, streptavidin(85878, Sigma-Aldrich)-coated, quartz slide, pretreated with NaCl imaging buffer for 30 min to minimize the oligomerization.

The surface-bound capsid proteins were illuminated using the evanescent field generated by the total internal reflection of a 532-nm laser. Fluorescence emission was collected through a 1.49 NA 100 × oil-immersion objective (Olympus), and passed through a filter to remove Rayleigh scattering. Acceptor and Donor emissions were separated using a dichroic mirror into green and red emissions and projected onto the EMCCD. The single-molecule time traces of around 100s were collected at a rate of 10 frames/second using Micro-Manager(µManager, https://micro-manager.org).

### Flow Cytometry

Trypsin digested cells were suspended, incubated with mAb 5E11 for 1.5 h at 37 °C, stained with FITC - goat anti-mouse IgG (ab6785, Abcam) for 1 h, and evaluated using a BD Accuri™ C6 Plus flow cytometer (BD Bioscience), and the ratio was analyzed using Flow Jo X 10.0.7 software(FlowJo, LLC, https://www.flowjo.com).

### High Performance Liquid Chromatography with Heparin-Sepharose

Purified capsids (100 µg) were loaded onto a 5-ml HiTrap™ Heparin-Sepharose HP Column (GE Healthcare), which then washed with Ca2+ and Mg2+-free PBS to remove unbound molecules and eluted with gradually linear increasing concentrations of NaCl. The original sample, flow through, and elution fractions were collected, diluted to same concentration, and then analyzed using SDS-PAGE and immunoblotting.

### Microscale Thermophoresis Assay

The Cy5-labeled capsid was incubated at a constant concentration of 20 nM with two-fold serial dilutions of heparin in MST-optimized buffer (50 mM Tris-HCl, pH 7.4, 150 mM NaCl, 1 mM MgCl2, 0.05% Tween-20). The mixtures were incubated for 15 min and added into glass capillaries and loaded into a Monolith NT.115 instrument (NanoTemper Technologies, Germany), the Kd values were determined using NanoTemper Analysis software (NanoTemper Technologies, https://nanotempertech.com).

### Construction of a PCV2 Infectious Clone and Virus Rescue

Site-specific mutagenesis to construct the 99R (SM) and 99R100K (DM) mutants was performed directly using PCR.

The linear genomes were extracted, cyclized using T4 DNA ligase (Takara), and transfected into PK15 cells with the jetPRIME® in vitro transfection reagent (CPT114vJ Polyplus-transfection). The transfected cells were cultured for 72 h and continuously passaged. The rescued samples were evaluated using indirect immunofluorescence assay and virus titers were calculated according to the Reed-Muench method.

### Analysis of smFRET Data

Using custom made IDL software (Exelis Visual Information Solutions, https://www.exelisvis.com), hundreds of dual-labeled single-molecule spots were picked from the images of an acceptor detection channel by alternating the 532 and 633 nm laser excitations. The intensities of Alexa 547 and Cy5 were analyzed by running the VBFRET software package script(The Gonzalez Laboratory, http://www.columbia.edu/cu/chemistry/groups/gonzalez/software) using a custom-made MATLAB algorithm (Mathworks, https://www.mathworks.com). The FRET efficiencies were calculated using the equation IA/ (IA + ID). All observed FRET data points were compiled into histograms using origin 9.0 software (Originlab, https://www.originlab.com/), and histograms were fitted using three Gaussian distributions. The occupancy of each Gaussian peak was calculated as the ratio of peak covering area.

### Statistical Analysis

All results are presented as means ± the standard deviation (SD). Significant differences between treated and control groups were analyzed using Student’s t test. The differences were considered significant and extremely significant at *P* values < 0.05 and < 0.01, respectively.

## Acknowledgments

This work was supported by the National Key Research & Development Program of China (2016YFD0500102, 2015BAD12B01) and the National Natural Science Foundation of China (grant numbers 31230072, 31402198).

## Author Contributions

J.Y.Z., J.R.L and J.Y.G. conceived the experiments. J.R.L and J.Y.G. prepared the samples and conducted most experiments. S.N.W and J.L. performed immunoblotting assays and constructed the infectious clone. J.W.Z., C.L. performed the purification of virus-like particles and observation of TEM. J.Y.Z. and J.R.L. analyzed and interpreted the data and wrote the manuscript.

## Declaration of Interests

The authors declare no competing interests.

---

**Figure S1.**
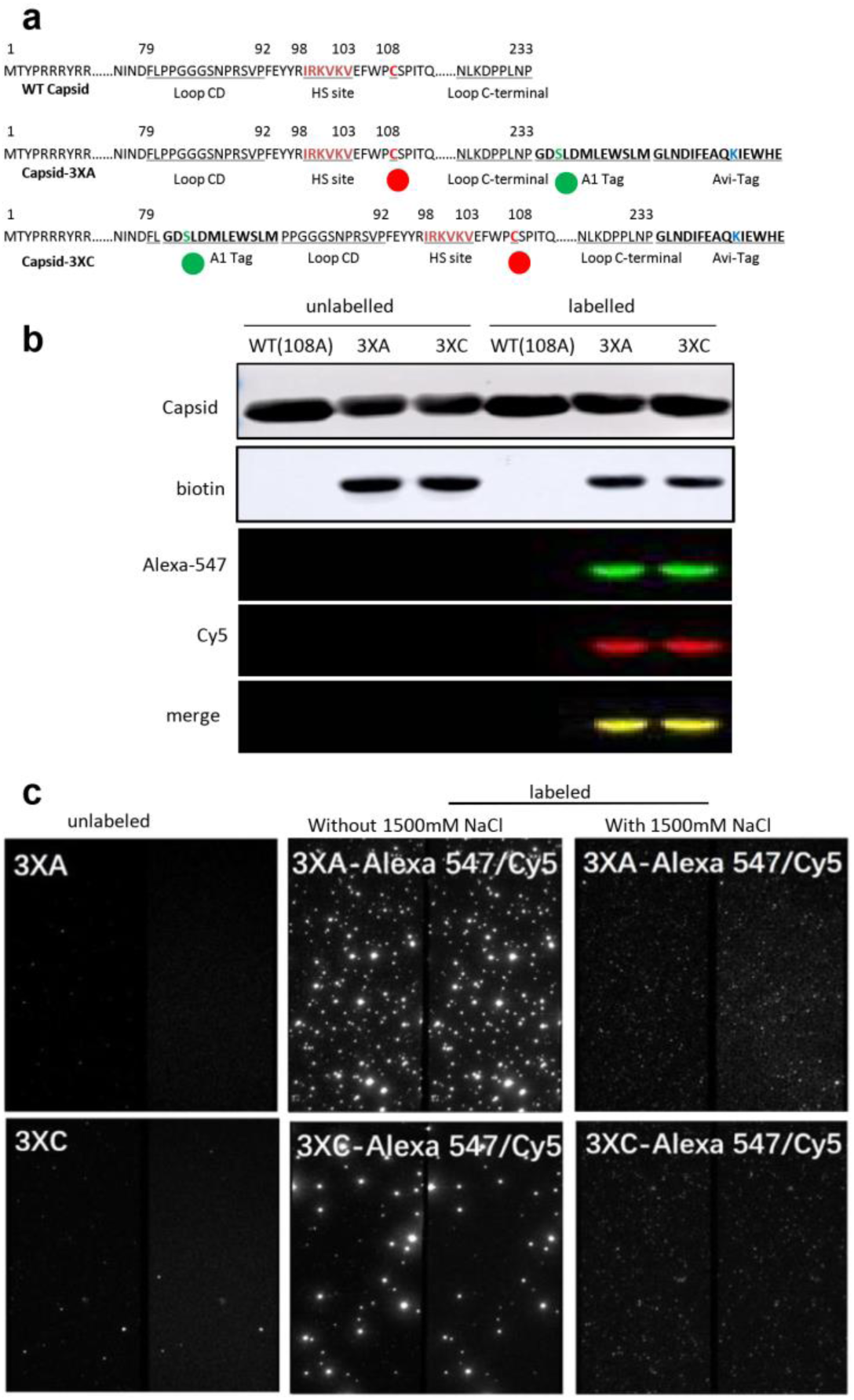
Strategy for Site-specific Attachment of the Donor & Acceptor Fluorophore Pair. (a) Schematic of the engineered capsids 3XA and 3XC, based on the WT capsid sequence of the PCV2 HZ0201 isolate. The putative heparin binding motif 98IRKVKV103 is highlighted in crimson. 108cysteine is highlighted in red and labeled by Cy5 (red ball). The A1-tag peptide (GDSLDMLEWSLM) was inserted at the C-terminus (3XA) or between the 80Leucine and 81Proline residues (3XC) to attached Alexa-547 (green ball). The Avi-tag peptide (GLNDIFEAQKIEWHE) was added at the C-terminus for the adhesion of D-biotin. (b) Western blotting assay of Cy5/Alexa 547 labeled capsid-3XA and capsid-3XC. Alexa-547 and Cy5 were excited by lasers at 520 nm and 630 nm, respectively. The conjugated biotin was identified using HRP-streptavidin. (c) Capsids were subjected to the fluorescent labeling reaction, surface immobilized, and imaged using TRIF. The donor and acceptor channels view of the passivated slides immobilized with unlabeled dCapsid 3XA or 3XC (left); with labeled dCapsid 3XA or 3XC (middle); or with labeled dCapsid 3XA or 3XC pretreated with 1500 mM NaCl imaging buffer for 30 min to eliminate oligomerization (right).

**Figure S2.**
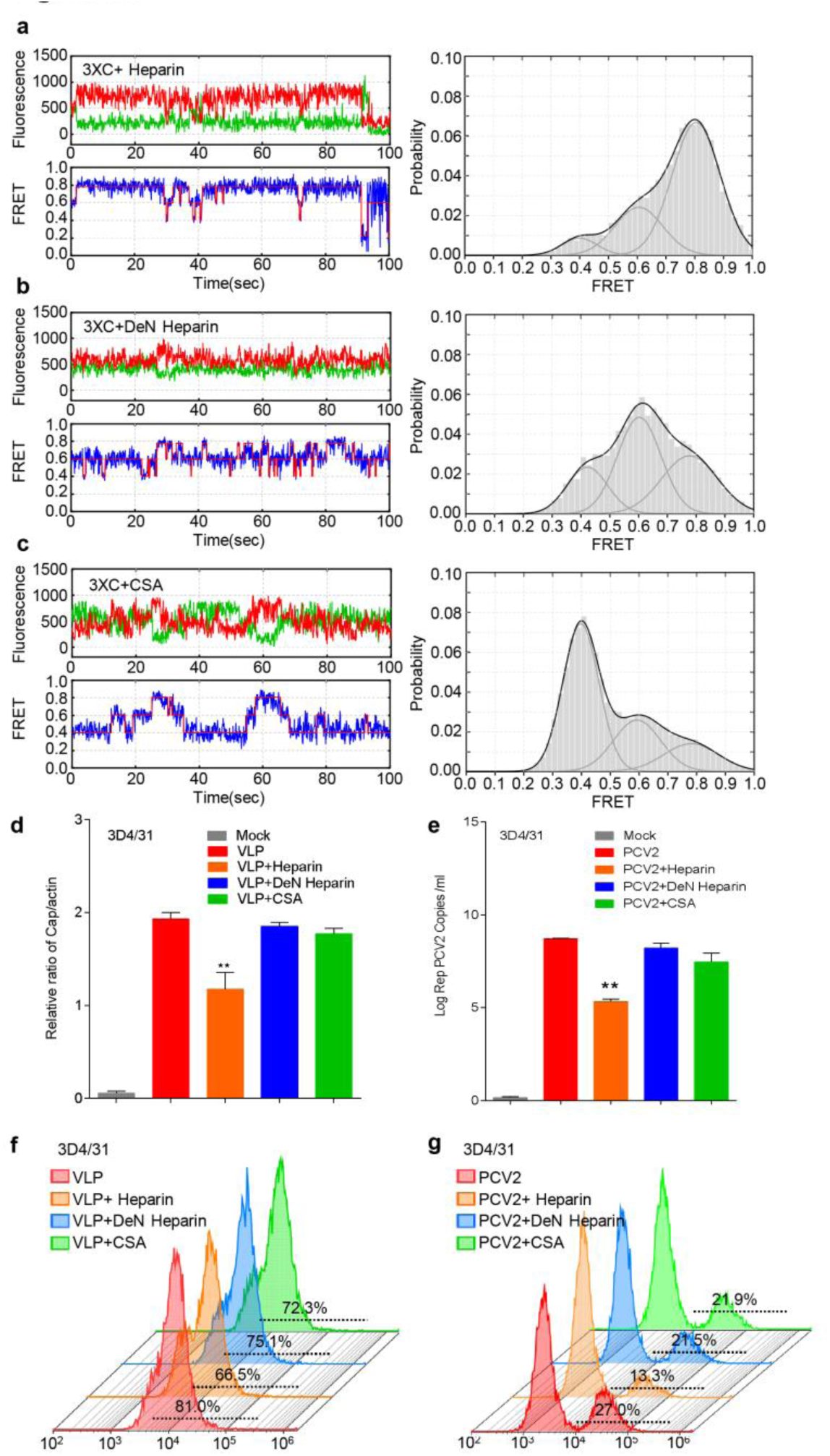
Interaction with Heparin Induced the High-FRET Conformation of Capsid. (a-c) Typical Capsid-3XC fluorescence time trace and the histogram distribution with exogenous soluble GAGs. (a), Capsid-3XC mixed with heparin. (b), Capsid-3XC mixed with De-N-sulfated acetylated heparin. (c), Capsid-3XC mixed with chondroitin sulfate A. The concentration of GAGs was fixed at 2,500 μg/ml. (d-g) Competitive binding assay of VLPs and PCV2 particles for 3D4/31 cells. The experimental protocol is identical to that detailed in Figure 3. (d), Western blotting analysis of VLPs bound to soluble GAGs. E, qPCR assay of PCV2 virions bound with soluble GAGs. (f), percentage of VLPs bound to 3D4/31 cells, as assessed using flow cytometry. (g), Percentage of PCV2 bound to 3D4/31 cells, as measured using flow cytometry.

**Figure S3.**
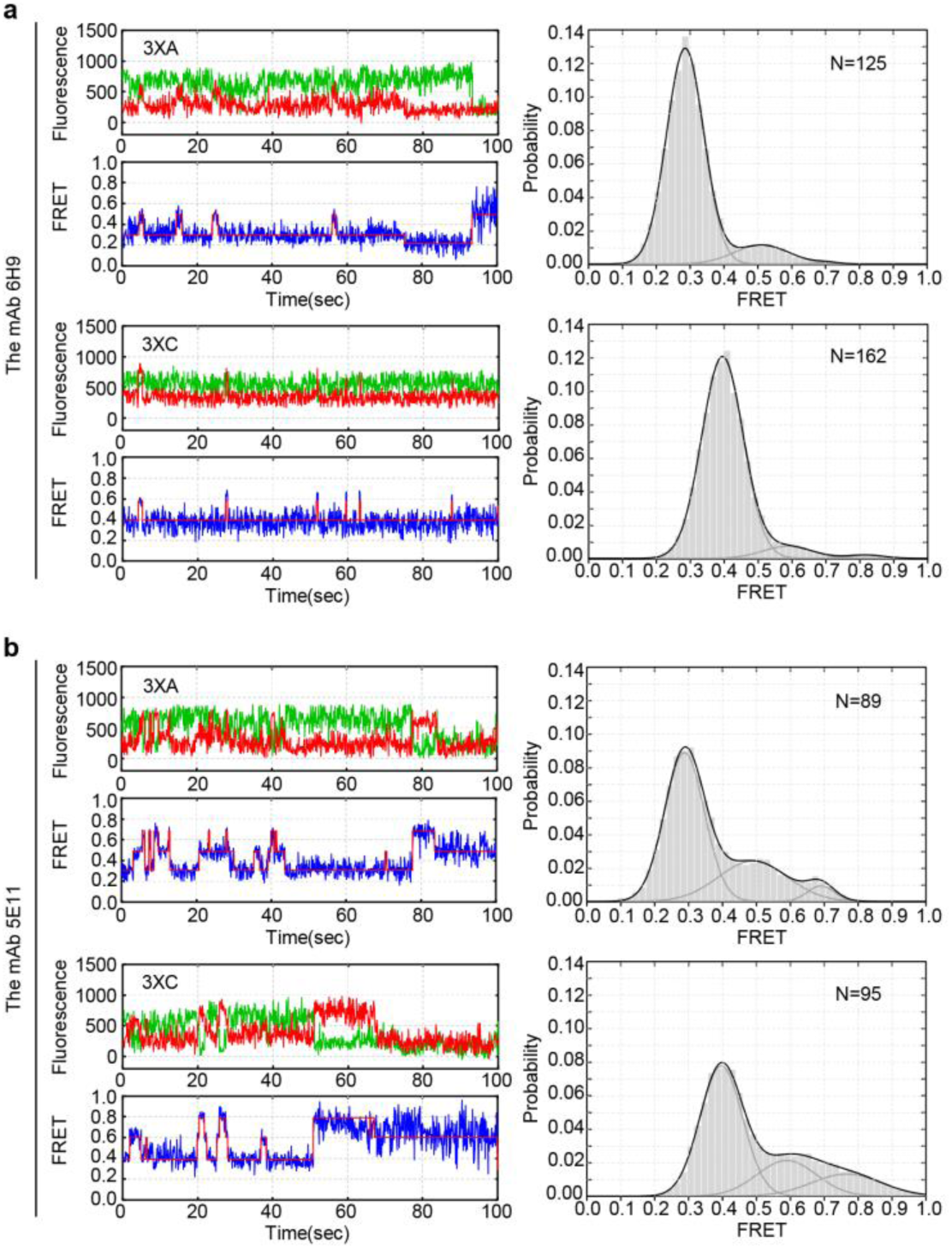
The Conformational Landscapes of Capsids Treated with Anti-Capsid Antibodies. The typical fluorescence time trace and histogram distribution of capsids 3XA and 3XC reacting with mAbs. (a), Capsids treated with neutralizing mAb 6H9 reveal an absolute predominance of the low-FRET state. (b), Capsids treated with non-neutralizing mAb 5E11 display a similar conformational distribution to free capsids (See Figure 2).

**Figure S4.**
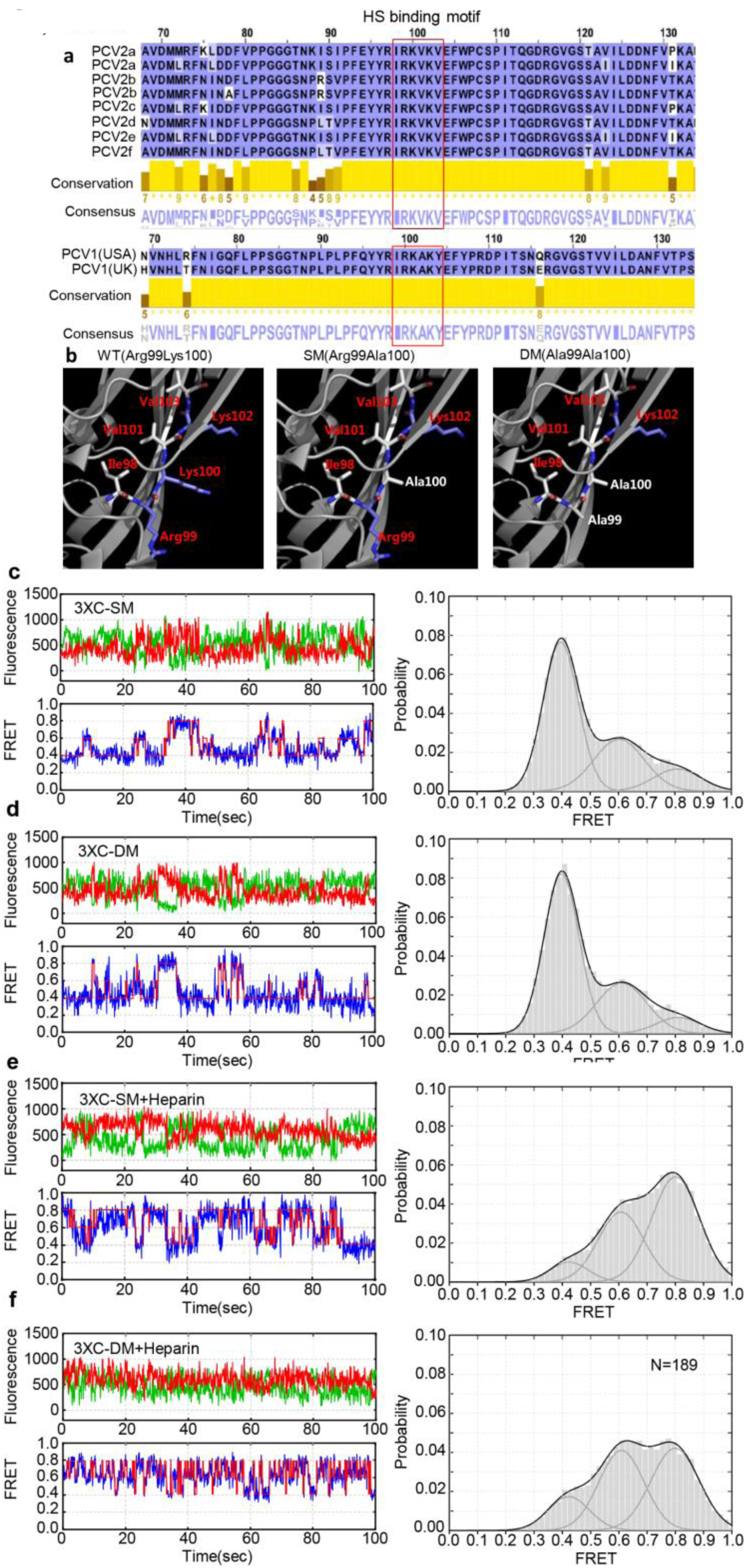
Deficiency of Positively Charged Amino Acids of the Binding Site Affects the Affinity to Heparin. (a) The conservation of the putative heparin sulfate binding motif in the capsid sequence. All the amino acid sequence data were obtained from GenBank. The location and sequence of the canonical putative heparin sulfate binding motif is highlighted using a red frame. (b) Schematic of the predicted local structure of the heparan binding site in the positively charged amino acid deficient mutants. The positively charged amino acids are labeled in blue and the uncharged amino acids are labeled in white. Left, the abundant side chain powered by the basic amino acids in capsid-WT. Middle, the side chain was reduced because of the K100A mutation in Capsid-SM. Right, the side chain was reduced because of the 99R100K to 99A100A mutations in Capsid-DM. This local structural prediction was performed on the basis of the revealed capsid structure (PDB:3R0R) and presented using the PyMOL software. (c-d) Typical smFRET trajectories and histogram distributions of Capsid-3XC-SM (c) and DM (d). Conformational features of the positively charged amino acids deficiency mutants are similar to those of free Capsid-WT. (e-f) Representative Capsid-3XC fluorescence time trace and the histogram distribution of Capsid-SM(e) and Capsid-DM(f) with added exogenous heparin.

